# Superior stimulation of female fecundity by subordinate males provides a mechanism for telegony

**DOI:** 10.1101/191510

**Authors:** Sonia Pascoal, Benjamin J. M. Jarrett, Emma Evans, Rebecca M. Kilner

## Abstract

When females mate promiscuously, sperm compete within females to fertilise the ova. In theory, a male can increase his success at siring offspring by inducing the female to lay more eggs, as well as by producing more competitive sperm. Here we report that the evolutionary consequences of fecundity stimulation extend beyond rival males, by experimentally uncovering effects on offspring. With experiments on the burying beetle *Nicrophorus vespilloides*, we show that smaller subordinate males are better able to stimulate female fecundity than larger, dominant males. Furthermore dominant males also benefit from the greater fecundity induced by smaller males, and so gain from the female’s earlier promiscuity just as predicted by theory. By inducing females to produce more offspring on a limited resource, smaller males cause each larva to be smaller, even those they do not sire themselves. Fecundity stimulation thus promotes the non-genetic inheritance of offspring body size, and provides a mechanism for telegony.

## Introduction

Sperm competition arises throughout the animal kingdom whenever females mate with more than one male (Parker 1970, 1998). Recent work has emphasised that a male’s success at sperm competition derives not only from his investment in high quality sperm but from his ability to manipulate female fecundity as well (e.g. Cameron et al. 2007, Parker & Pizzari 2010 Alonzo & Pizzari 2010, Perry et al. 2013). By inducing a female to produce more eggs, through courtship feeding or nuptial gifts or through direct physiological manipulation via components of his ejaculate, a male can potentially increase the number of offspring he sires-even if his share of paternity remains relatively low.

In theory, the extent to which males should invest in simulating female fecundity depends on the male’s mating role, that is whether his mating behaviour consistently places him at an advantage or disadvantage in sperm competition (Cameron et al. 2007, Parker & Pizzari 2010, Alonzo & Pizzari 2010). A male’s mating role might be conferred on him by his social status. For example, dominant males consistently occupy the favoured role through their ability to mate more frequently, and last, with the female (e.g. Lemaitre et al. 2012). Holding a particular mating role changes the payoffs derived from investing in fecundity stimulation relative to other strategies for enhancing fertilisation success (Cameron et al. 2007 Alonzo & Pizzari 2010, Lemaitre et al 2012). It also sets up producer-scrounger dynamics between rival males, in which a later mating male can potentially parasitise any previous investment in female fecundity stimulation by earlier mates of the same female (Alonzo & Pizzari 2010). However, whether socially dominant and subordinate males differ in the extent to which they invest in fecundity stimulation is not yet known.

A major consequence of male fecundity stimulation, which has thus far been relatively neglected, is the effect on the offspring (Crean et al. 2016). A few studies have considered the direct effect that components of the male’s ejaculate might have on the offspring’s phenotype (e.g. Garcia-Gonzalez & Dowling 2015, Crean et al. 2016). Yet there is a very well-characterised trade-off between offspring number and offspring size, from diverse species across the animal kingdom (Stearns 1992, Rollinson & Rowe 2015). Thus fecundity stimulation could influence the offspring’s phenotype simply through reduced nourishment of each egg (e.g. Nager et al. 2000) or through increased competition among offspring for limited resources during offspring development (e.g. Mock & Parker 1997). Through this simple mechanism, a female’s previous mates could influence her offspring’s phenotype even if they sire none of her offspring, a phenomenon known as telegony. But whether this mechanism exists in reality is so far unknown.

Here we determine whether males of different social status differ in the extent of their fecundity stimulation and whether the stimulation of female fecundity alone is sufficient to change the offspring’s phenotype. Our experiments focus on burying beetles, *Nicrophorus vespilloides*. Burying beetles breed on a small dead vertebrate, like a mouse, which they require to provision their larvae (Scott 1998). There is competition for this scarce resource and disputes are settled by fighting within each sex. The outcome determines an individual’s social status during that breeding event (Müller et al. 1990, Eggert & Müller 1992, Pettinger et al. 2011). The winners are usually the largest male and female (Scott 1998, Hopwood et al 2016a) and they become the dominant pair on the carcass. They gain most reproductive success on the carcass, and stay to defend and care for the larvae (Eggert & Müller 1997). Defeated, usually smaller, individuals become subordinate satellites, Subordinate males gain reproductive success by sneaking matings with the dominant and other females (e.g. Müller et al. 2007). Females become subordinate co-breeders (Eggert & Müller 1992) or intraspecific brood parasites (Müller et al. 1990), depending on the size of the carcass. Regardless of their social status, females are highly promiscuous (Müller & Eggert 1989, Müller et al. 2007, House et al. 2007, 2009). Furthermore, previous work has shown that female fecundity is increased by multiple mating in a dose-dependent way (House et al. 2009).

## Results

We analysed the effect of a male’s social status on fecundity stimulation by using body size as a proxy for dominant (= large) or subordinate (= small) status. We began by phenotypically engineering males and females of different sizes, within the natural range, by varying the extent of their nourishment while larvae (see Methods). Males were either ‘Large’ or ‘Small’, while females were of intermediate size (see Methods). Upon reaching sexual maturity, these males and females were then divided into four treatment groups. Females were allowed to mate for an equal time period with two different males in succession, generating four treatments in all: a Large male followed by a Small male (LS) and a Small male followed by a Large male (SL), a Large male followed by another Large male (LL) and a Small male followed by another Small male (SS) (see Methods). Upon removal of the second male, the female was given a carcass of standard size for raising offspring. We counted the number of eggs she laid, and the number and mass of larvae she produced. Paternity of the offspring was assigned using microsatellite markers (Pascoal & Kilner 2017, see Methods).

As shown in previous work (Müller & Eggert 1989; Müller et al. 2007), we found that the last male to mate with the female typically obtained most paternity. However, we also found that the P2 values differed between males in the two different size treatments (estimated effect = 0.59 ± 0.59, z = 3.35, P = 0.001, Figure 1). For Small males, P2 was roughly 50% whereas for Large males P2 was approximately 75% (Figure 1), regardless of the size of the first male to mate with the female. Overall, we found P2 was considerably lower than reported in previous studies on *N. vespilloides*, which used sterile males or a phenotypic marker to assign paternity (63% of all offspring vs c. 90% from previous work (e.g. Eggert & Müller 1989, House et al. 2007, 2008)).

**Figure 1.**
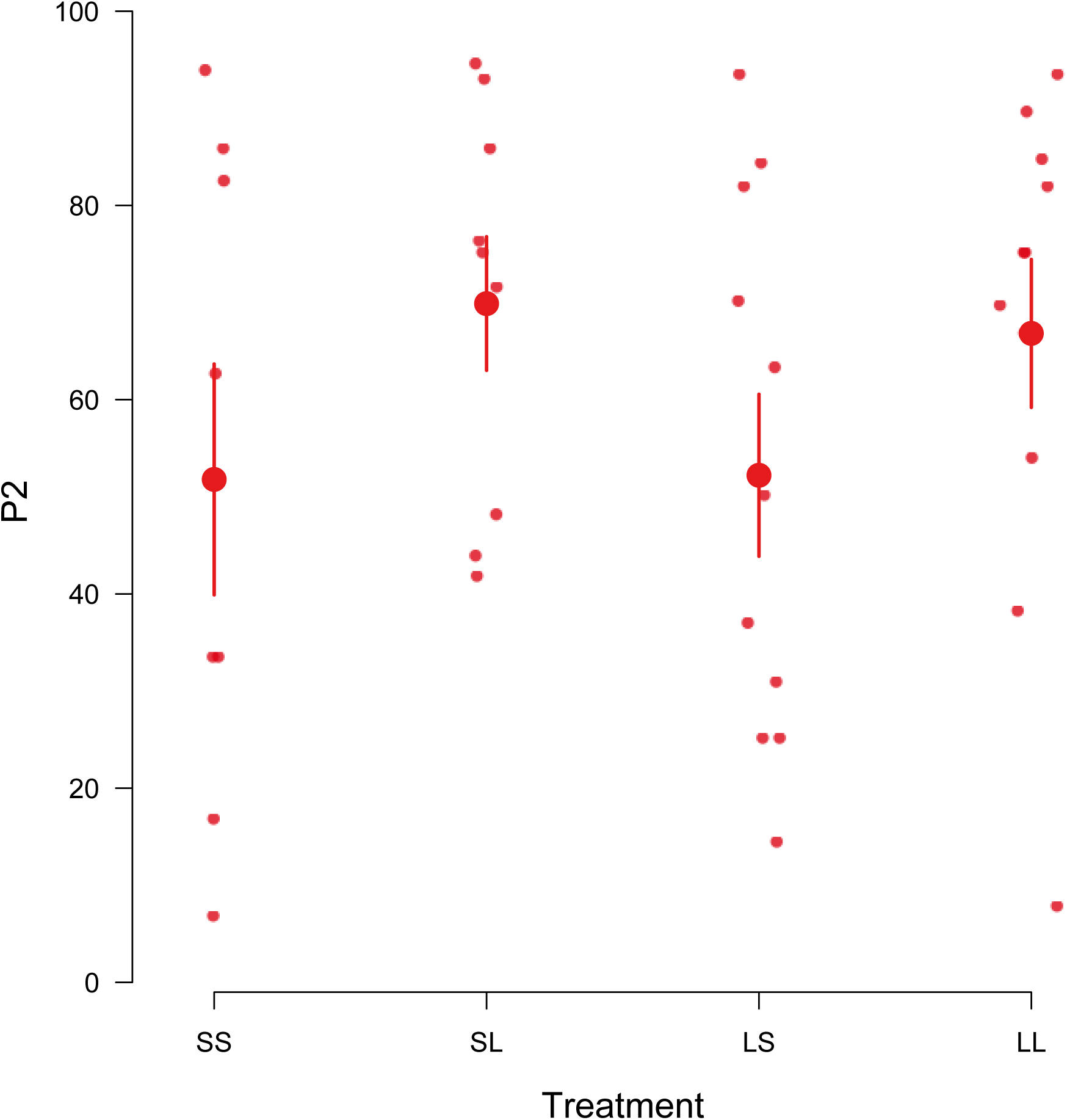
P2 scores (measured as % of the brood sired by the second male to mate with the female) for males in each of the four treatments in the experiment. Each female was mated twice, with the following treatments: SS = Small male followed by a Small male; SL = Small male followed by a Large male; LS = Large male followed by a Small male; LL = Large male followed by a Large male. Each datapoint represents a brood. Large points are the treatment means with standard errors.

There was no significant interaction between the size of the first and second males that influenced P2 values (estimated effect = 0.45 ± 0.35, z = 1.27, P = 0.20), nor did the size of the first male influence the proportion of the brood that he sired (estimated effect =0.13 ± 0.18, z = 0.70, P = 0.48, Fig. 1), Carcass size (estimated effect = 0.23 ± 0.17, z = 1.40, P =0.16), and female size (estimated effect =0.49 ± 0.33, z = 1.50, P = 0.13) were each unrelated to P2 values. We cannot infer from our data why Large males obtained larger P2 scores. It is possible that they produced more competitive sperm, or ejaculates that better promoted fertilisation success (Perry et al. 2013). It is just as possible that females simply mated more frequently with Large second males than with Small second males (cf Moya-Marano & Fox 2006).

We found that Small males were more effective at stimulating female fecundity than were Large males (Figure 2a). When Small males were first to mate, females then laid significantly more eggs than when Large males were first to mate, (z = 2.64, P = 0.008,Figure 2a). Carcass mass independently and positively influenced clutch size, (z = 4.10, P < 0.001) There was no interaction between the size of the first male and the size of the second male on clutch size, (z = 1.26, P = 0.21), and nor did size of the second male influence clutch size (z = -1.40, P = 0.16). These differences in fecundity persisted until larvae dispersed away from the carcass to pupate (Figure 2b). Broods were larger when Small males mated first than when Large males mated first, (z = 2.96, P = 0.003, Fig. 2b). Carcass size did not explain variation in brood size (z = 0.88, P = 0.38). There was no interaction between the first and second males in determining brood size, (z = 0.43, P = 0.67) nor did second male size have any effect (z = -0.87, P = 0.38).

**Figure 2.**
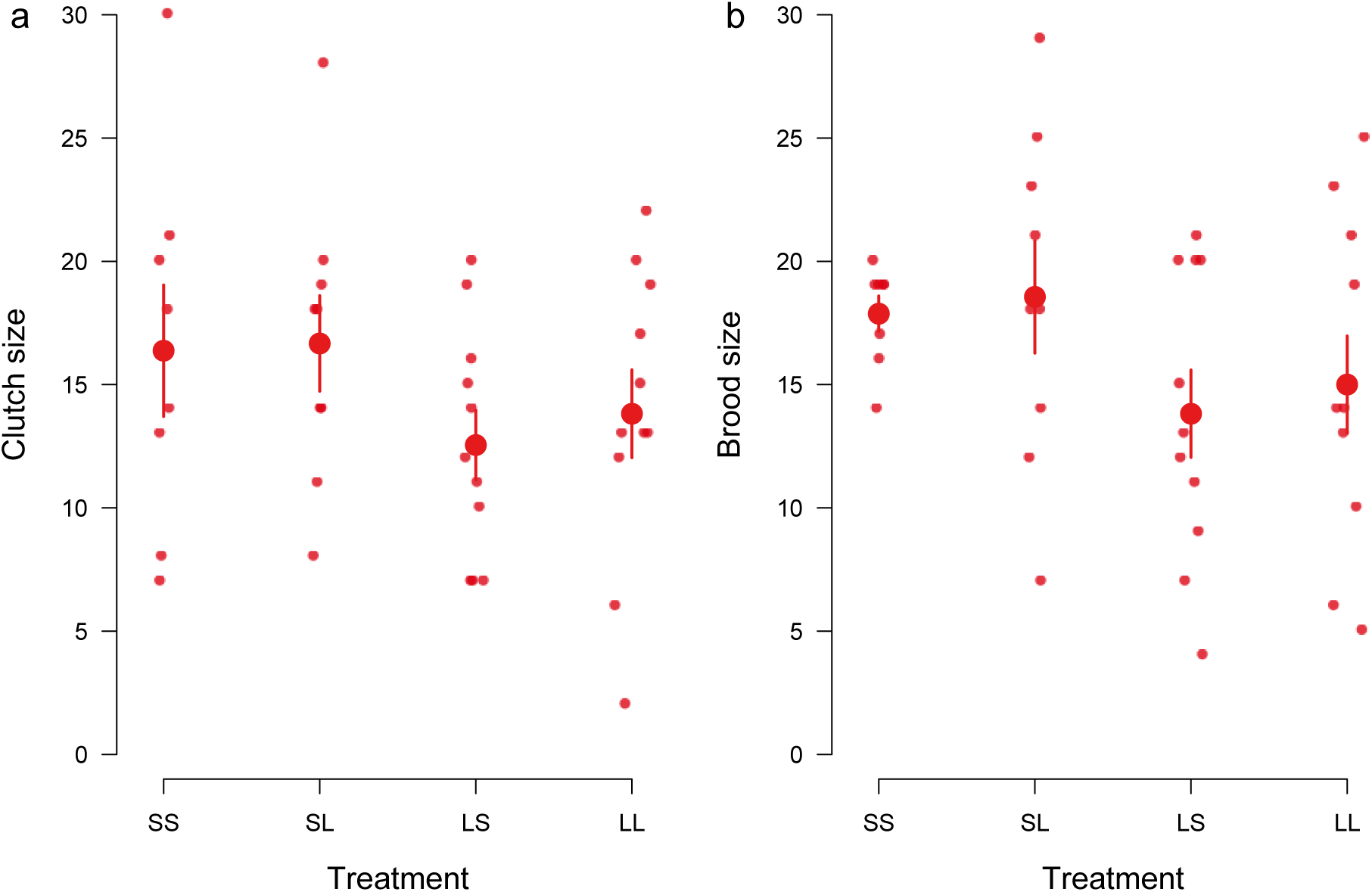
The effect of male size and mating order on a) clutch size and b) brood size. Each female was mated twice, with the following treatments: SS = Small male followed by a Small male; SL = Small male followed by a Large male; LS = Large male followed by a Small male; LL = Large male followed by a Large male. Each datapoint represents a brood. Large points are the treatment means with standard errors

The stimulation of female fecundity is a public good (Cameron et al 2007, Alonzo & Pizzarri 2010) and therefore potentially of benefit to all the males that mate with a female. We investigated whether both males benefited from the increase in clutch size induced when Small males mated first. We found some evidence that Small males could enhance their reproductive success through fecundity stimulation. Small males that mated first sired more larvae than Large males that mated first - but this was only true when the second male was Small (Figure 3a, z = 2.60, P = 0.009). When the second male was Large, his greater P2 score overwhelmed any advantage the Small male might have gained through fecundity stimulation. We also found evidence that Large males mating second benefitted from the increased clutch size stimulation by the Small male mating first. They produced more offspring than Small males mating second after a different Small male (Figure 3b, z = 2.86, P = 0.02).They also tended to produce more offspring than the Large males mating second after another Large male, though not significantly so (Figure 3b, estimated effect (Figure 3b, estimated effect = 0.28 ± 0.14, z = 2.10, P = 0.15).

**Figure 3.**
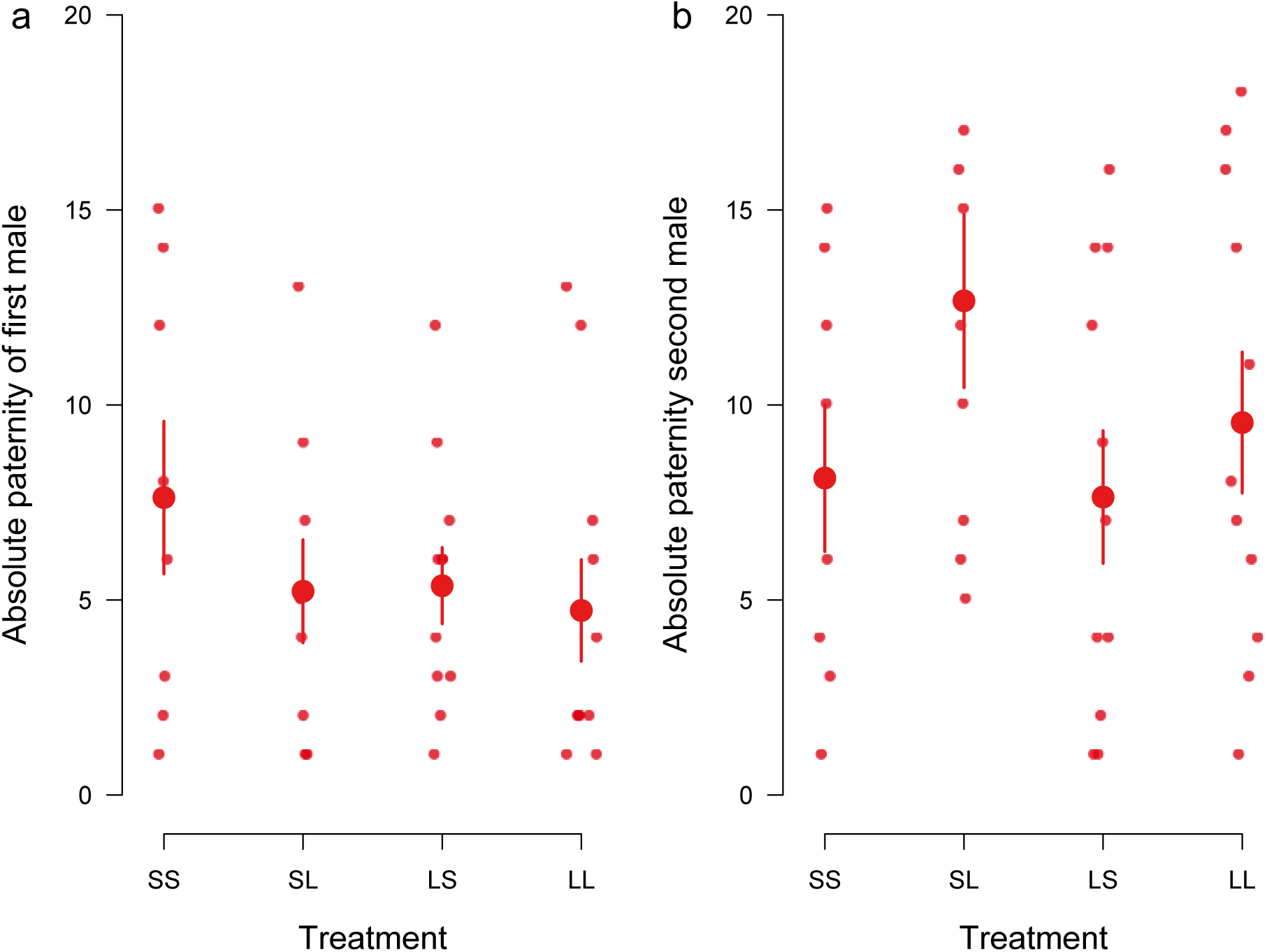
Effect of fecundity stimulation on number of offspring produced by a) first-mating males andb)second-mating males. Each female was mated twice, with the following treatments: SS = Small male followed by a Small male; SL = Small male followed by a Large male; LS = Large male followed by a Small male; LL = Large male followed by a Large male. Each datapoint represents a brood. Large points are the treatment means with standard errors.

Although our experimental design deliberately minimized variation in female size, it was impossible to eliminate all variation experimentally. Since female size can independently account for variation in clutch size (e.g. Schrader et al 2016), it might mask more subtle effects of any male-induced effects on her fecundity. To control for the possibility, we next incorporated female size into analyses of fecundity stimulation. This exposed effects of the second male on clutch size (Figure 4a). Furthermore, we found that Small second males were especially effective at inducing larger females to lay more eggs (Figure 4a, estimated effect of second Small male = 1.35, se = 0.34, z = 3.98, P < 0.001). However, larger females were more likely to lay fewer eggs when second males were Large (Figure 4a). These results show that Small males were more effective than Large males at stimulating female fecundity, even when they mated second. They also reveal size-related variation in the female’s response to fecundity stimulation, with clutch size declining with female size when second males were Large, but rising with female size when second males were Small. When we repeated these analyses using brood size at dispersal, rather than clutch size as the measure of fecundity, the effects persisted in a similar direction but were no longer as great in magnitude, nor were they significant (Figure 4b, estimated effect of second Small male = 0.40, se = 0.32, z = 1.28, P = 0.20).

**Figure 4.**
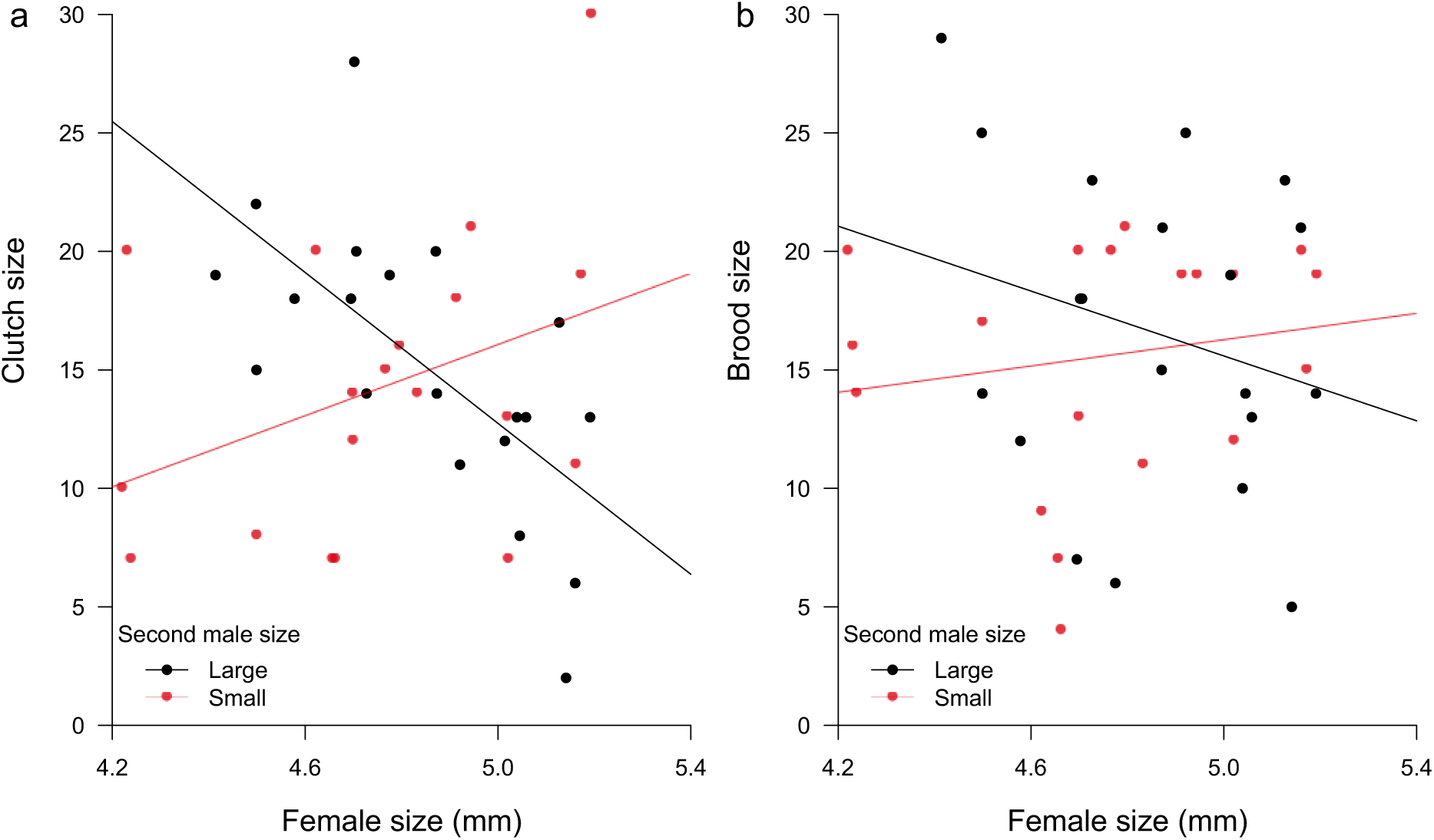
The effect of female size and the size of the second male on a) clutch size and b) brood size. The red datapoints and line indicate the second male was small and the black points and line indicate the second male was large. Linear regression lines are plotted

In our final analysis, we investigated the effects of fecundity stimulation by males on offspring size. Each carcass bears finite resources for nourishing the brood, and previous work on burying beetles has identified a pronounced trade-off between brood size and offspring size (e.g. Schrader et al. 2015). We found the same trade-off here, with a similarly steep negative gradient irrespective of whether the first male to mate with the female was Small or Large (Figure 5a, t = 0.42, P = 0.68). Since Small males induce females to produce more offspring (Figure 2) they should also cause females to produce smaller offspring, irrespective of whether they have sired the offspring. Comparing average larval mass across the four mating treatments we found that when Small males mated first, larvae were indeed smaller at dispersal than when Large males mated first (Figure 5b, estimated effect=-0.02,se = 0.01, t = -2.03, P = 0.049).

**Figure 5.**
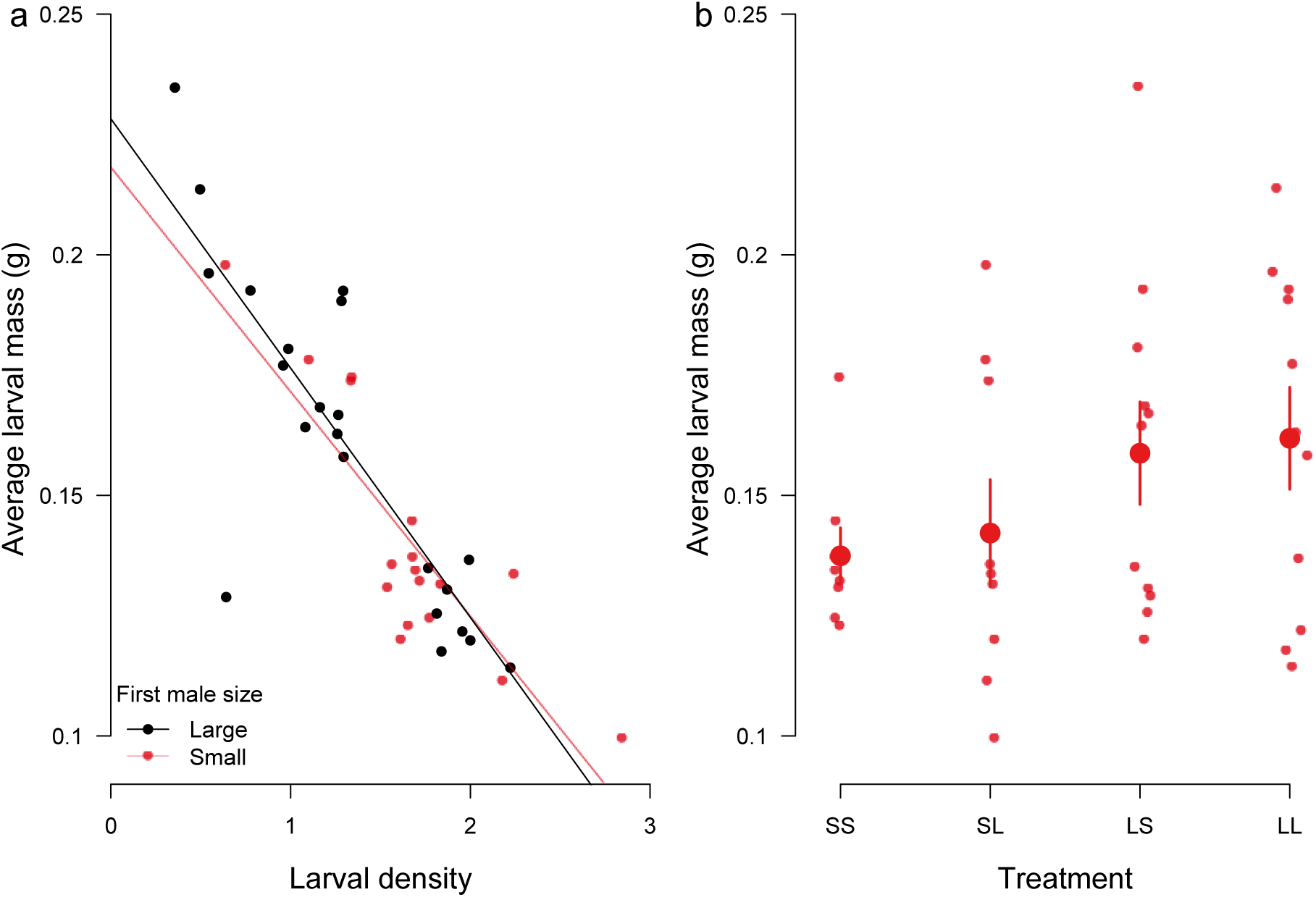
(a) The trade-off between larval density and larval mass at dispersal when first males to mate are large (red line and red datapoints) or small (black line and black datapoints).. Linear regression lines are plotted. (b). Average larval mass across the four mating treatments. Each female was mated twice, with the following treatments: SS = Small male followed by a Small male; SL = Small male followed by a Large male; LS = Large male followed by a Small male; LL = Large male followed by a Large male. Each datapoint represents a brood. Large points are the treatment means with standard errors.

Thus the fecundity stimulating effect of Small first males causes a reduction in offspring size. Small males induce the production of smaller offspring, regardless of whether the Small male is their sire. Fecundity stimulation therefore provides a general non-genetic mechanism for the cross-generational transmission of body size and a simple mechanism for telegony, where offspring inherit characteristics of their mother’s previous mates (Crean et al. 2014). Nevertheless, the effects of the Small males on larval size at dispersal were weaker than their effects on clutch size. This suggests that females may be able to counteract any negative effects on offspring size of over-producing larvae, and that these measures occur between egg-laying and larval dispersal. A likely counter-measure, known to happen in burying beetles, is partial filial cannibalism of first instar larvae (Bartlett 1988).

## Discussion

Our results reveal that Small males are more effective than Large males at stimulating female fecundity, but do not indicate how this is achieved. Since there is no courtship behaviour in burying beetles, nor the presentation of any nuptial gifts, nor any pheromonal displays when beetles are in close proximity, we suggest these effects are most likely due to differences in ejaculate composition. Detailed analyses of *Drosophila* and *Tribolium* ejaculates have found that they contain a multitude of proteins which alter female physiology in diverse ways (Sirot et al. 2011, Yamane et al. 2014, Bayram et al. 2017, Wigby et al. 2016), and that smaller *Drosophila* allocate more proteins from their accessory gland to their ejaculate than do larger males (Wigby et al. 2016). Previous work on *Tribolium* beetles further suggests that the same proteins that promote fecundity stimulation might also reduce egg fertilisation success (Yamane et al. 2014). This could explain why we found Small burying beetle males to be both better at fecundity stimulation and to have relatively low P2 scores. However, all this remains to be investigated since nothing is yet known about the constituents of burying beetle ejaculates. An alternative interpretation is that females, and not males, caused the effects we found. They may have detected the size of their mates and differentially produced eggs accordingly. We cannot rule out this suggestion. But nor can we explain how it would be adaptive for a female to allow Small, but not Large, males to induce her to lay more eggs.

However, we can explain why it would be adaptive for males of different sizes to differ in the extent to which they stimulate female fecundity. The adaptive reasoning stems from differences between Small and Large males in their capacity to win fights over a carcass (e.g. Otronen et al. 1988, Hopwood et al. 2016a, Müller et al. 2007). A Small male is lucky if he secures carcass ownership outright and is unlikely to be as fortunate again in future breeding attempts. Fecundity stimulation can help him capitalise on his good fortune by pursuing a near semelparous reproductive strategy. A more likely scenario is that he becomes a satellite subordinate, and reliant on sneaking fertilisations with a dominant female to gain fitness. Our data suggest that here too, fecundity stimulation is potentially adaptive because it increases the chance that he will fertilise some eggs. Larger males are less dependent on stimulating female fecundity because they are more likely to win contests for a carcass, and consequently better able to monopolise matings with the dominant female (e.g. Otronen et al. 1988, Hopwood et al. 2016a, Müller et al. 2007, Pettinger et al 2011). Nevertheless, and just as predicted by theory (Alonzo & Pizzari 2010), we have shown that they can profit from the increased fecundity stimulated by female’s earlier promiscuity with other males, providing they sire a high proportion of the brood.

From the female’s perspective, it is presumably beneficial to outsource fecundity stimulation to the male, at least to some extent (Alonzo & Pizzari 2010). Nevertheless, we found evidence to suggest that females vary in their response to fecundity stimulation in a complex way, according to their size, and the size of the second male they mated with (Figure 4), even though we deliberately minimised variation in female size experimentally. Since a female’s social status also varies with size in burying beetles (Muller et al. 1990, Muller & Eggert 1992), it raises the previously unexplored possibility that a female’s response to fecundity stimulation might vary adaptively, according to the mating strategy associated with her social status. This will determine the benefits she stands to gain from fecundity stimulation relative to the costs she incurs. In cooperative breeders with helpers and a high level of reproductive skew, for example, it may be beneficial for a dominant female to be susceptible to fecundity stimulation because then she can gain extra offspring without paying all the costs of raising them. The same reasoning could apply to subordinate interspecific brood parasites. By contrast, any female that is likely to pay a sub-optimally high cost for producing more young will benefit by resisting fecundity stimulation (Lessells 2006). It would be interesting to explore these possibilities in future theoretical and empirical work.

In summary, competition for a carcass breeding resource in burying beetles causes large males to become dominant and smaller males to be subordinate. Dominants and subordinates then pursue contrasting mating strategies, which intensify sperm competition. We have shown that smaller males can enhance their success in the competition for fertilisations by more effectively stimulating female fecundity. We have also shown that large males can profit from the fecundity stimulating actions of their female’s previous mates. Finally, we have demonstrated that the greater stimulation of female fecundity by smaller males causes the production of smaller offspring, a finding that solves the puzzle of evolutionary stasis in burying beetle body size (Hopwood et al. 2016b). The novel insight from our experiment is that there are opposing effects on body size of competition before and after mating. Competition for a carcass persistently selects for larger individuals. But sperm competition favours smaller, subordinate males that more effectively stimulate female fecundity and this causes the production of smaller individuals. The variance in body size might be increased as a result of these two opposing effects, but the mean will remain unchanged.

## Material and Methods

### Maintenance of the beetle population

The *N. vespilloides* population used in the experiment was established in 2014 from wild beetles caught from three sites (Gamlingay Woods, Waresley Woods, and Byron’s Pool) in Cambridgeshire, UK. Wild caught beetles were added every two weeks from June to October each year to ensure the population was outbred. Adult beetles were fed twice a week with raw beef mince and kept individually in plastic boxes (12x8x6cm) filled with moist soil. Adults were sexually mature at two weeks post-eclosion. They were bred at 2-3 weeks post-eclosion by placing a male and female together in a breeding box (17x12x6cm) lined with soil and furnished with a mouse carcass (8-14g). The breeding boxes were left in a dark cupboard to simulate the underground conditions where breeding would naturally occur. Eight days after pairing, the larvae were ready to disperse from the carcass, at which point they are collected, counted, and weighed. They were then placed into cells (2x2x2cm) in an eclosion box (10x10x2cm) filled with peat until they were fully developed adults, three weeks after dispersal. Both individual boxes and eclosion boxes were kept out in the laboratory which was maintained on a 16L:8D hour light cycle at 21°C.

Adult beetle size was determined by measuring the widest part of the pronotum, a commonly used and accurate proxy for adult size in beetles (e.g. Painting & Holwell 2013; Tomkins et al. 2005). To do this, beetles were photographed individually using a mounted digital camera and a custom MATLAB script was used to determine pronotum width (version 8.5.0 2015).

### Experimental design

The experiment consisted of two steps: 1) generating beetles of different sizes, and then 2) measuring the effect of i) male size on fecundity stimulation and ii) fecundity stimulation on offspring size.

#### Step 1: Manipulation of Beetle Size

Three groups of experimental subjects were created in this step: intermediate-sized females, large males, and small males. To achieve this, a male and a female burying beetle were placed in a breeding box, one-third filled with moist soil. The mated pairs were 2-3 weeks old, were not siblings and were both virgins. To breed intermediate sized females, mating pairs were given an 8-14g freshly defrosted mouse carcass. After eclosion, the beetles were sexed: the females were retained and the males were discarded.

To manipulate male size, mating pairs were given a mouse carcass weighing 21-26g. Five days after pairing, half the larvae were removed from the carcass to eclosion boxes. This early removal, before natural dispersal, prevented carcass consumption and so yielded Small individuals (from methods used by Steiger 2013). The larvae that remained on the carcass, and were now destined to be Large, were removed 8 days after pairing (which is when larvae typically disperse from the carcass) and transferred to eclosion boxes. After eclosion, individuals from both these treatments were sexed. The males were kept and the females were discarded.

The pronotum width of beetles from all three groups of retained offspring was measured at eclosion. Males of intermediate size were discarded to ensure that there was no region of overlap between the Large and Small males. Large males were therefore significantly larger than Small males (t-test: t_76_ = 26.6, P < 0.0001). Large and Small females were also discarded to ensure that any differences detected between treatments could be attributable to the greater variation in male size, and mating sequence. The remaining experimental beetles were then left for two weeks to reach sexual maturity. The pronotum width of all the experimental beetles fell within the range observed in natural populations of this species (range of beetles found in the wild: 3.10 - 6.01 mm; range in this experiment:3.32-5.90 mm, Sun et al., unpubl data, Kilner et al., 2015).

#### Step 2: Fecundity stimulation by males, and effects on females and offspring

In burying beetles the dominant male on the carcass holds the favoured role in sperm competition because he can monopolise matings with the female over a prolonged period and just prior to egg production (Pettinger et al. 2011). These males are usually also larger and therefore in better condition. Satellite males are disfavoured by both the relative lack of mating opportunities (Pettinger et al. 2011) and by being smaller. Our experiment was designed to break up the usual correlation between mating opportunities and male size, so that we could more confidently attribute a male’s ability to gain paternity and stimulate fecundity to male size alone. Furthermore, the procedure for mating the beetles was designed to maximise the exposure of the female to each male, so that any effects we detected on fecundity stimulation and paternity were more likely to be explained by events after mating rather than opportunities for mating. (Note that there is no courtship in this species). Thus we are not attempting to estimate the likely share of paternity in the wild by recreating natural conditions for mating but rather to test specifically for evidence that males of different social status by virtue of their size (dominant = Large; subordinate = Small) differ in the extent to which they can stimulate a female’s fecundity.

To achieve this, males and females were divided into four treatment groups. Females were each mated successively with the two types of males in a fully crossed design, comprising: a Large male followed by a Small male (LS) and a Small male followed by a Large male (SL), a Large male followed by another Large male (LL) and a Small male followed by another Small male (SS). Within each experimental trio, the first male (M1), the second male (M2) and the female (F), were all unrelated. Each trio comprised adults that derived from a unique combination of broods, to prevent any confounding effects that might be attributable to the family of origin.

The mating procedure began when we placed a virgin female in a breeding box with the first male for 24h. The first male was then exchanged with the second male who remained with the female for a further 24h. When the second male was removed, the female was given a 10-12g mouse carcass (mean=10.95g, SD=0.59) to breed upon. By removing males after mating, we eliminated any potential confounding effects of paternal care. The breeding boxes were filled with only 1cm of soil, making it possible to count the number of eggs each female laid. At dispersal, eight days later, the larvae were weighed individually to within 0.001g. After eclosion, offspring pronotum width was measured.

Parents and offspring from the successful breeding attempts (N=63 total; SS=13, LL=18, SL=15, LS=17) were preserved in absolute ethanol for genetic analysis.

### DNA extractions and parentage analysis

We used microsatellites to assign paternity. Total genomic DNA (n=1005; 204 parents of known sex and 801 offspring) was individually extracted from beetle heads using the DNeasy Tissue Kit (Qiagen) following the manufacturer’s instructions. For parentage analysis, up to 9 previously developed polymorphic microsatellite markers (Pascoal & Kilner 2017) were used (Table S1). All individuals were genotyped for 5 markers (mix1) and, when necessary (n=359), for additional 4 markers (mix2) to increase confidence of parentage assignment. Microsatellite amplification and multiplexing was performed as described in Pascoal & Kilner (2017). Briefly, two microsatellite multiplexes were amplified using the Qiagen Multiplex PCR kit. Genotyping was performed on an ABI 3730 instrument at the Edinburgh Genomics Institute Sanger Sequencing Centre with GeneScan 500 LIZ (Applied Biosystems) as internal size standard. Alleles were scored and checked using Peak Scanner v.1.0 (Applied Biosystems) and parentage analysis was performed using CERVUS (Kalinowski et al. 2007). The number of alleles scored in all tested individuals (n = 1005) for the 9 polymorphic microsatellite markers ranged between 7 and 15 (Table S1). For comparison with previous studies, we calculated P2 scores as the share of paternity gained by male mating second with the female (Table S2).

### Data analysis

#### Effect of male size on P2 and fecundity stimulation

We used R (version 3.3.2) for all statistical analyses. The dataset we analysed included only the families where both males had sired at least one offspring each. In this way, we could be confident that both males had successfully mated with the female and that each male tested was reproductively competent.

Since there was no overlap in male size between the Large and Small treatments, we coded for male treatment in our analyses by using a two level factor. In all analyses, the interaction between the first male (M1) and the second male (M2) was included at first, and then removed if non-significant. Block was always included as a random term in the global model, but was always removed if it did not improve the fit of the model.

The proportion of offspring sired by the second male to mate (ie the P2 score, given by the number offspring sired by the second male in relation to the total number of offspring produced) was analysed with the cbind function in a glm with a binomial error structure. To measure fecundity stimulation, we analysed the effect of the male on clutch size and brood size, using a generalised linear model (glm) with the Poisson error term and log link function. The size of the carcass was added as covariate in the model. Residuals were plotted and diagnostic plots were examined for all models ensuring all analyses were appropriate. For measures of fitness, we included only the absolute number of offspring sired as opposed to the proportion of paternity attained. The number of offspring sired, therefore, was analysed with the interaction of the two male treatments in a glm with a poisson error distribution and log link function.

To understand which males benefitted from fecundity stimulation, we analysed the absolute number of offspring sired by each male using a glm with poisson error structure and log link function. To compare the numbers of offspring sired between treatments, the four treatments were treated as an independent factor with four categories and differences between treatments were analysed using post-hoc comparisons.

#### Controlling for female size on the extent of fecundity stimulation

Here we examined the interaction between the size of the female the size of her first and second mate. If this three-way interaction was non-significant, it was dropped from the model. We used a glm with poisson error structure and log link function to analyse these effects on clutch size and brood size. The model was simplified until the minimal model remained.

#### Effect of fecundity stimulation on offspring size

The average size of the larvae for each brood was analysed using a linear model which included the interaction between the size of the first male and size of the second male. This interaction term was dropped from the model if non-significant. As the effect of male size on offspring size is mediated through changes in brood size, we did not fit larval density as a term in the model. Terms were removed until the minimal model was found.

## ACKNOWLEDGEMENTS

This project was supported by a Consolidator’s Grant from the European Research Council (310785 BaldwinianBeetles) and by a Wolfson Merit Award from the Royal Society, each to RMK. Emma Evans was funded by the Balfour-Browne Fund, administered by the Department of Zoology at the University of Cambridge, and by Pembroke College, Cambridge. We are indebted to Sue Aspinall and Chris Swannack for maintaining the beetle populations in the laboratory. We also thank Goran Arnqvist, Tracey Chapman, Jennifer Perry and Stuart Wigby for stimulating and insightful discussions.

## SUPPLEMENTARY MATERIAL

**Table S1.**
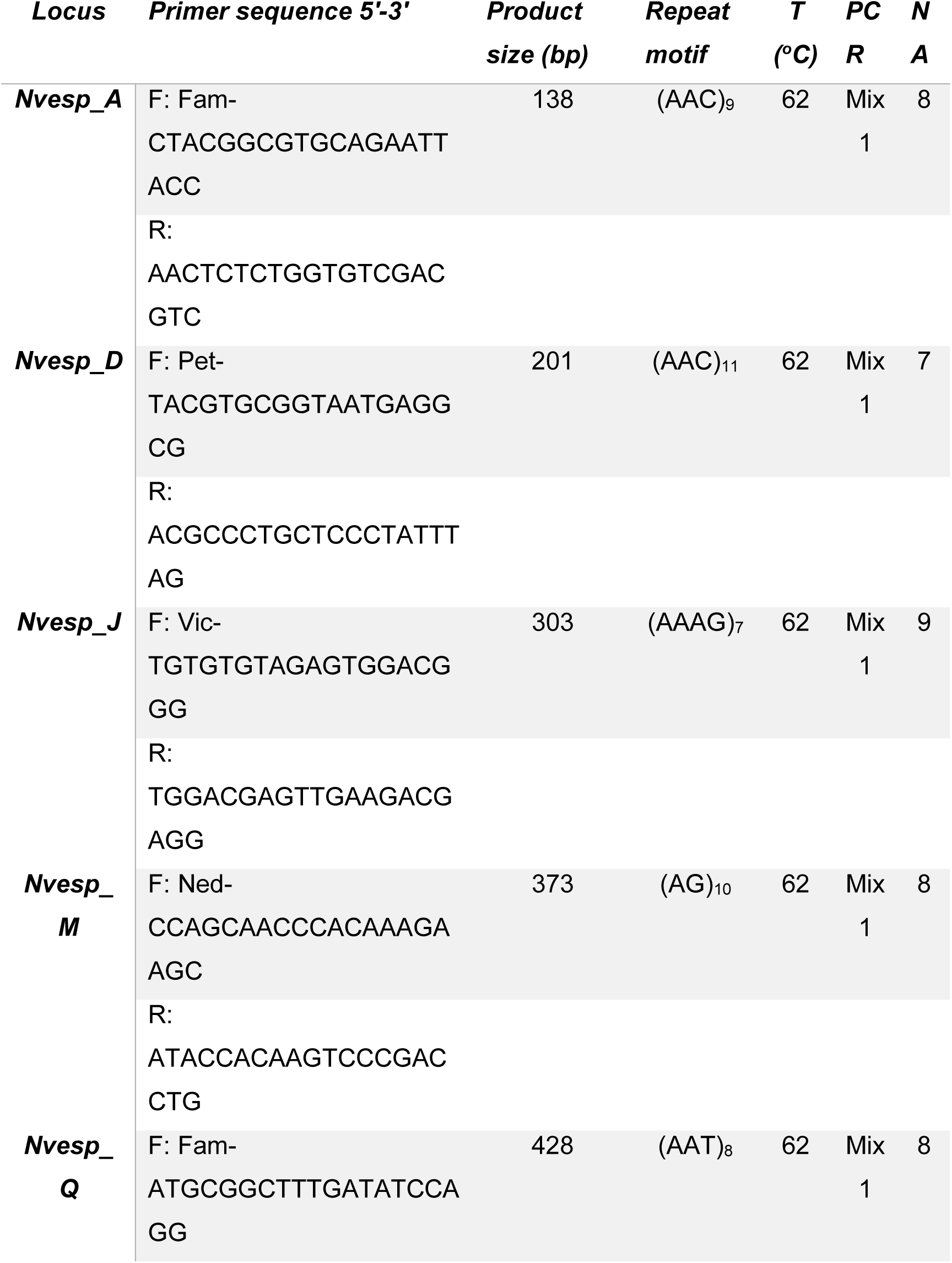
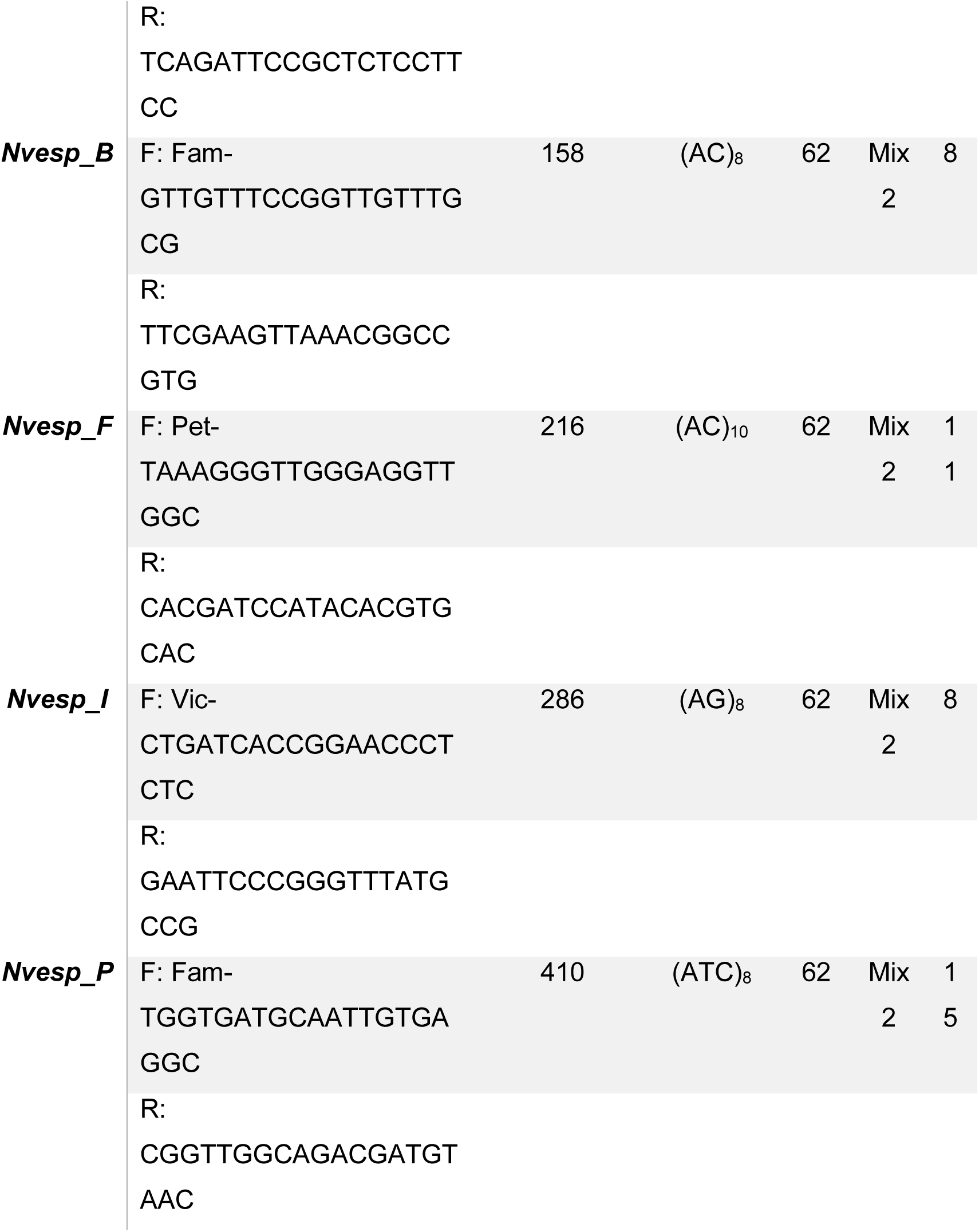
Details of the *Nicrophorus vespilloides* microsatellite markers used for parentage analysis. NA: number of alleles per locus.

**Table S2:** Summary of the parentage assignment analysis per treatment; SS = Small male followed by a Small male; SL = Small male followed by a Large male; LS = Large male followed by a Small male; LL = Large male followed by a Large male. #: number of individuals.

|  | *Offspring #* | *M1 #* | *M2 #* | *% M2* |
| --- | --- | --- | --- | --- |
| **SS03** | 15 | 14 | 1 | 7 |
| **SS05** | 8 | 0 | 8 | 100 |
| **SS06** | 16 | 1 | 15 | 94 |
| **SS07** | 4 | 4 | 0 | 0 |
| **SS09** | 17 | 3 | 14 | 82 |
| **SS12** | 18 | 12 | 6 | 33 |
| **SS13** | 12 | 8 | 4 | 33 |
| **SS14** | 18 | 15 | 3 | 17 |
| **SS16** | 14 | 2 | 12 | 86 |
| **SS17** | 16 | 0 | 16 | 100 |
| **SS18** | 9 | 9 | 0 | 0 |
| **SS19** | 6 | 6 | 0 | 0 |
| **SS20** | 16 | 6 | 10 | 63 |
| **SL04** | 15 | 0 | 15 | 100 |
| **SL05** | 7 | 1 | 6 | 86 |
| **SL06** | 14 | 0 | 14 | 100 |
| **SL08** | 16 | 9 | 7 | 44 |
| **SL10** | 6 | 0 | 6 | 100 |
| **SL11** | 21 | 5 | 16 | 76 |
| **SL12** | 25 | 13 | 12 | 48 |
| **SL13** | 13 | 0 | 13 | 100 |
| **SL14** | 28 | 2 | 26 | 93 |
| **SL15** | 18 | 1 | 17 | 94 |
| **SL17** | 20 | 5 | 15 | 75 |
| **SL19** | 12 | 7 | 5 | 42 |
| **SL20** | 14 | 4 | 10 | 71 |
| **LL03** | 5 | 0 | 5 | 100 |
| **LL05** | 16 | 0 | 16 | 100 |
| **LL06** | 19 | 2 | 17 | 89 |
| **LL07** | 6 | 2 | 4 | 67 |
| **LL08** | 8 | 2 | 6 | 75 |
| **LL09** | 13 | 12 | 1 | 8 |
| **LL10** | 21 | 13 | 8 | 38 |
| **LL11** | 2 | 0 | 2 | 100 |
| **LL12** | 23 | 7 | 16 | 70 |
| **LL13** | 13 | 2 | 11 | 85 |
| **LL14** | 3 | 3 | 0 | 0 |
| **LL15** | 7 | 0 | 7 | 100 |
| **LL16** | 15 | 1 | 14 | 93 |
| **LL17** | 22 | 4 | 18 | 82 |
| **LL18** | 13 | 6 | 7 | 54 |
| **LL19** | 4 | 1 | 3 | 75 |
| **LL20** | 21 | 0 | 21 | 100 |
| **LL21** | 2 | 0 | 2 | 100 |
| **LS01** | 8 | 4 | 4 | 50 |
| **LS02** | 13 | 13 | 0 | 0 |
| **LS03** | 20 | 6 | 14 | 70 |
| **LS04** | 13 | 9 | 4 | 3185 |
| **LS06** | 8 | 6 | 2 | 25 |
| **LS07** | 19 | 12 | 7 | 37 |
| **LS08** | 4 | 3 | 1 | 25 |
| **LS09** | 11 | 2 | 9 | 82 |
| **LS10** | 19 | 7 | 12 | 63 |
| **LS11** | 21 | 0 | 21 | 100 |
| **LS13** | 19 | 3 | 16 | 84 |
| **LS14** | 15 | 0 | 15 | 100 |
| **LS15** | 16 | 16 | 0 | 0 |
| **LS18** | 15 | 1 | 14 | 93 |
| **LS19** | 7 | 6 | 1 | 14 |
| **LS20** | 2 | 2 | 0 | 0 |
| **AvgSS** | 13 | 6 | 7 | 47 |
| **AvgSL** | 16 | 4 | 12 | 79 |
| **AvgLL** | 12 | 3 | 9 | 74 |
| **AvgLS** | 13 | 6 | 8 | 48 |
| **TotalSS** | 169 | 80 | 89 | 53 |
| **TotalSL** | 209 | 47 | 162 | 78 |
| **TotalLL** | 213 | 55 | 158 | 74 |
| **TotalLS** | 210 | 90 | 120 | 57 |

